# Rational and computation-assisted engineering of a compact and efficient StaCas9 genome editor

**DOI:** 10.64898/2026.07.21.739771

**Authors:** Miaomiao Li, Wenni You, Yuwen Tian, Jingtong Liu, Shengzhou Wang, Jinzhong Lin, Yongming Wang

**Affiliations:** Center for Medical Research and Innovation, Shanghai Pudong Hospital, Fudan University Pudong Medical Center, School of Life Sciences, Fudan University, Shanghai, 200438, China; Shanghai Engineering Research Center of Industrial Microorganisms, Shanghai, 200438, China

## Abstract

Type II-A CRISPR-Cas9 nucleases are widely used for genome editing, yet their functional diversity and therapeutic potential remain incompletely explored. In this study, we systematically analyzed natural variation in PAM recognition among SaCas9 orthologs and identified StaCas9 as a compact and efficient nuclease recognizing an NNG PAM. Structural and sequence analyses revealed that S983 within the PAM-interacting domain contributes to the PAM preference of StaCas9. StaCas9 enabled efficient genome editing across multiple endogenous loci in human cells and achieved high-efficiency disruption of the therapeutically relevant PCSK9 gene, supporting its potential for gene therapy. Furthermore, guided by structural insights, we significantly enhanced the editing activity of StaCas9. Together, these results expand our understanding of PAM recognition in type II-A Cas9 nucleases and establish StaCas9 as a high-performance genome-editing tool for therapeutic applications.

## Introduction

The advent of CRISPR–Cas9 technology has revolutionized genome engineering by enabling highly programmable and efficient DNA manipulation ^1, 2^. CRISPR–Cas9 systems are classified into four subtypes (type II-A, II-B, II-C, and II-D), among which type II-A and type II-C together account for more than 96% of all Cas9 orthologs identified in nature and have been most widely exploited for genome editing applications ^3–5^. Notably, type II-A Cas9 nucleases generally exhibit intrinsically higher DNA cleavage activities than type II-C Cas9, largely attributable to their superior DNA unwinding capability ^6, 7^. Accordingly, the two most extensively used Cas9 nucleases to date, SpCas9 and SaCas9, both belong to the type II-A subtype ^1, 8^.

SpCas9 remains the most broadly applied Cas9 owing to its high editing efficiency and simple NGG PAM requirement ^9, 10^. However, its large coding sequence (4,104 bp) precludes packaging into a single adeno-associated virus (AAV) vector together with sgRNA and regulatory elements, thereby limiting its in vivo therapeutic utility. To overcome this limitation, Ran et al. reported in 2015 a compact Cas9 nuclease from Staphylococcus aureus (SaCas9, 3,159 bp), representing a small type II-A Cas9 well suited for AAV-mediated delivery ^8, 11^. SaCas9 has demonstrated robust genome editing activity in mammalian systems and has been explored in multiple disease contexts, including sensory disorders and prion diseases ^12, 13^.

Despite these advantages, the stringent NNGRRT PAM requirement of SaCas9 substantially restricts its targetable genomic space. To expand the editing scope of compact type II-A Cas9s, several SaCas9 orthologs have been identified and engineered, including SauriCas9 and SlugCas9, both recognizing an NNGG PAM^14, 15^, as well as SchCas9, which recognizes an NNGR PAM ^16^. Nevertheless, even with these advances, large fractions of the genome remain inaccessible, highlighting a persistent unmet need for compact Cas9 nucleases with relaxed PAM constraints.

The ever-expanding genomic databases offer a powerful opportunity to overcome the intrinsic PAM constraints of compact Cas9 nucleases. By systematically mining NCBI genomic repositories, we identified StaCas9, a previously underexplored SaCas9-related ortholog that recognizes a minimal NNG PAM, thereby substantially expanding target site accessibility. StaCas9 exhibits efficient genome editing activity in mammalian cells. Furthermore, through structure-guided engineering, we developed an enhanced variant, enStaCas9, which displays markedly improved editing efficiency.

## Results

### Identification of active SaCas9 orthologs for mammalian genome editing

To discover compact Cas9 nucleases with expanded PAM recognition, we performed a systematic homology-based search of the NCBI genomic databases using SaCas9 as a reference. Guided by prior structural insights indicating that SaCas9 residue N985 plays a critical role in PAM recognition by forming a hydrogen bond with the fourth PAM nucleotide ^17^, we focused on SaCas9-related orthologs exhibiting divergence at this position. The residues corresponding to N985 are very conserved among previously studied SaCas9 orthologs (Fig. S1). We identified StaCas9 from *Staphylococcus* sp.17KM0847, a previously uncharacterized ortholog harboring a serine substitution (S983) corresponding to SaCas9 N985. Structural modeling suggested that SaCas9 N985 forms a hydrogen bond with the nucleotide at the SaCas9 PAM position 4, while StaCas9 S983 forms a hydrogen bond with K1014 instead of PAM (Fig. S2). These data indicate that StaCas9 may recognize a relaxed PAM distinct from previously characterized Cas9 nucleases. In addition, we identified three SaCas9 orthologs (HloCas9, SciCas9, and Sha4Cas9) with a smaller size due to 11 aa deletions in the PI domain (Fig. S1). Smaller Cas9 is suitable for AAV delivery, so we also included these three Cas9 nucleases (Table 1).

**Table 1.**
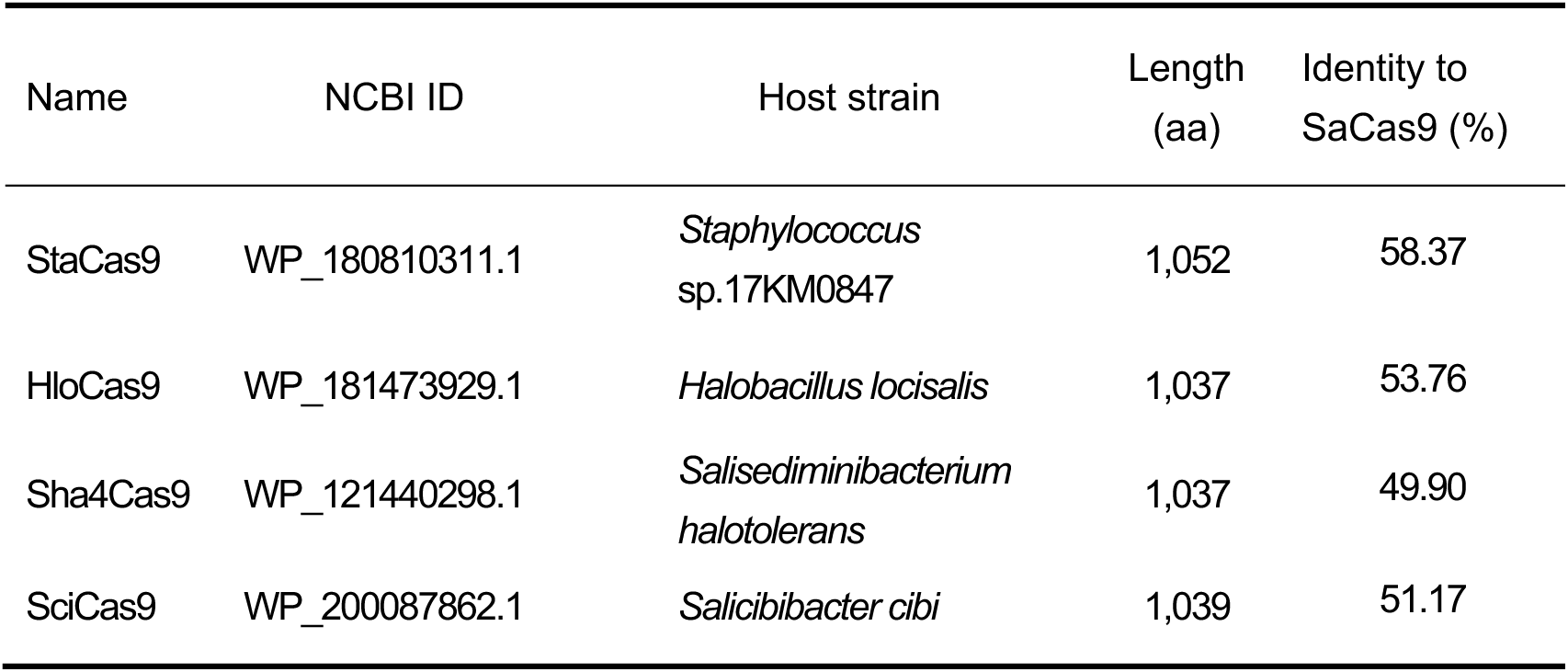
Four SaCas9 orthologs selected from the NCBI database.

Genomic locus analysis revealed that the StaCas9 locus encodes, in order, a CRISPR array, a tracrRNA (predicted based on complementarity to the repeat sequence), Cas9, Cas1, Cas2, and Csn (Fig. S3A). In contrast, the remaining three Cas9 loci encode the same set of elements but with the CRISPR array located downstream of the Cas genes. Sequence alignment of the CRISPR repeat and tracrRNA sequences revealed moderate conservation at the 5′ end, whereas the 3′ regions showed little conservation (Fig. S3B–C).

Based on these sequence features, we designed single-guide RNA (sgRNA) scaffolds for each ortholog by fusing the 3′ end of a truncated direct repeat to the 5′ end of the corresponding tracrRNA, including the full-length tail, using a 4-nt linker (Fig. S4A). Secondary structure prediction indicated that the StaCas9 sgRNA forms three stem– loop structures, similar to SaCas9, whereas the sgRNAs of the remaining three Cas9 orthologs form four stem–loop structures (Fig. S4B).

We next assessed the genome-editing activities of the identified Cas9 orthologs using a GFP-based functional reporter system ^18^. In this assay, a synthetic target cassette, comprising a 7-bp randomized sequence downstream of a protospacer, was embedded within the GFP coding region, resulting in a disrupted open reading frame (Fig. 1A). Genome editing–induced insertions or deletions (indels) at the target site can stochastically restore the correct reading frame, thereby reactivating GFP expression in edited cells (Fig. 1B).

**Figure 1.**
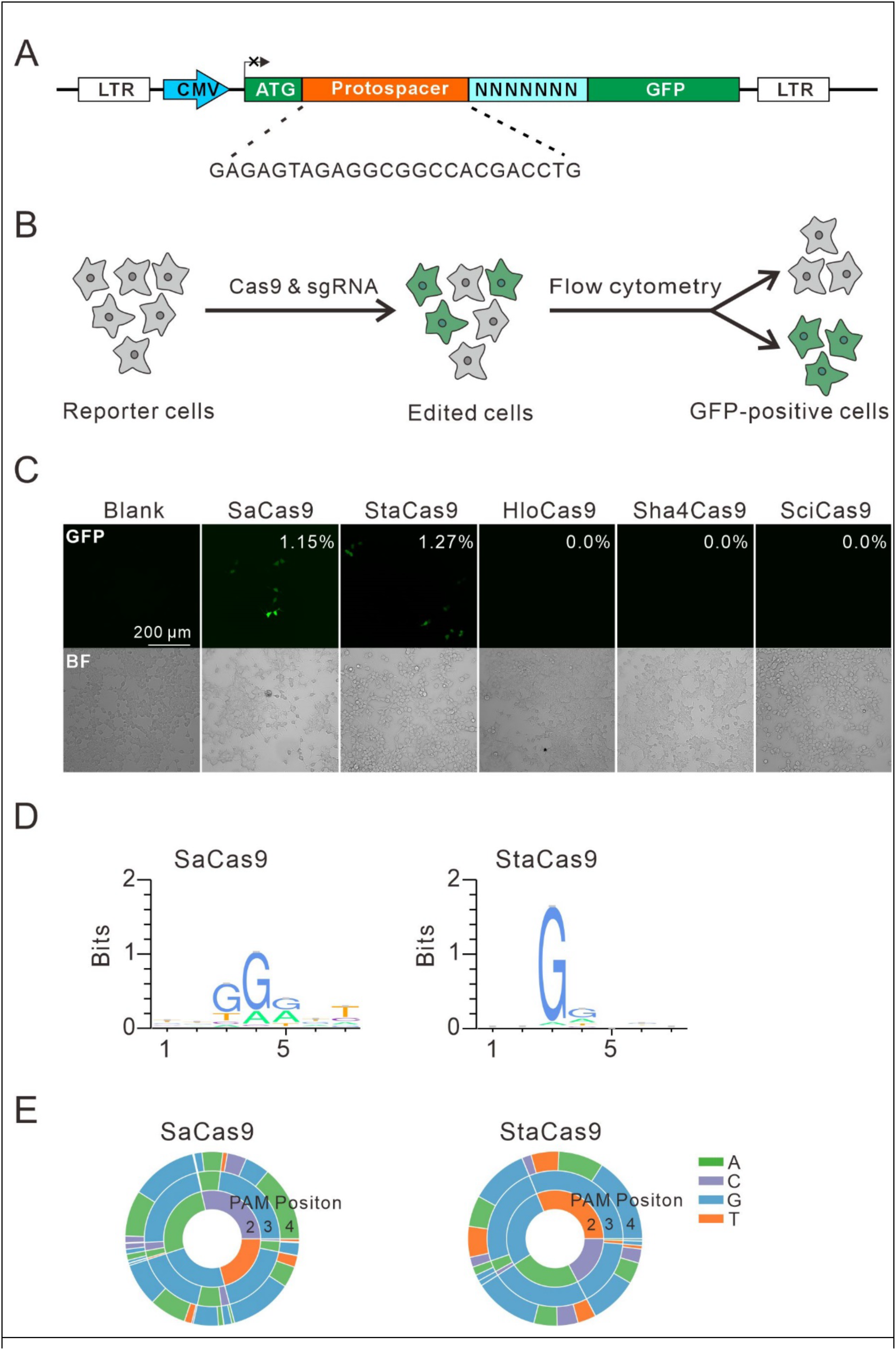
Functional screening of Cas9 ortholog activities using a GFP-activation reporter. **(A)** Schematic of the GFP-activation reporter. A target cassette consisting of a protospacer preceded by a 7-bp randomized sequence is inserted immediately downstream of the ATG start codon, disrupting the GFP open reading frame. The reporter library was stably integrated into HEK293T cells. **(B)** Workflow of the Cas9 activity screen. Reporter cells were co-transfected with Cas9 and sgRNA expression plasmids. Editing-induced insertions or deletions at the target site can restore the GFP reading frame in a subset of cells, resulting in GFP-positive signals. **(C)** Representative fluorescence images showing GFP activation induced by SaCas9 and StaCas9. **(D)** Sequence logos derived from deep sequencing of edited target sites, illustrating the PAM preferences of SaCas9 and StaCas9. **(E)** PAM wheel representations summarizing nucleotide enrichment patterns within the PAM regions based on deep sequencing analysis.

The reporter construct was stably integrated into HEK293T cells via lentiviral transduction. Each human-codon–optimized Cas9 ortholog was synthesized and cloned into a mammalian expression vector. Upon co-expression of each Cas9 ortholog with the SaCas9 sgRNA scaffold, robust GFP-positive populations were observed for both SaCas9, which served as a reference nuclease, and StaCas9, indicating that StaCas9 is catalytically active in mammalian cells (Fig. 1C). In contrast, no GFP activation was detected for HloCas9, Sha4Cas9, or SciCas9 under the same conditions.

To further assess sgRNA compatibility, each Cas9 ortholog was co-expressed with its corresponding sgRNA scaffold. Under these conditions, GFP-positive cells were detected only for StaCas9, whereas the remaining Cas9 orthologs failed to induce detectable editing activity (Fig. S5). Since the SaCas9 sgRNA scaffold and StaCas9 sgRNA scaffold displayed comparable activity, the SaCas9 sgRNA scaffold was used in the following study.

To define the PAM preferences of each nuclease, GFP-positive cells were sorted, and the edited target regions were subjected to high-throughput sequencing. Sequence enrichment analysis of the recovered target sites, visualized using WebLogo and PAM wheel representations, confirmed that SaCas9 exhibits a strong preference for the canonical NNGRRT PAM, in agreement with a previous report (Fig. 1D–E) ^8^. By contrast, StaCas9 displayed a markedly relaxed PAM profile, with a predominant NNG requirement and a modest enrichment for G or A at the fourth PAM position (Fig. 1D–E). Together, these data establish StaCas9 as a functional Cas9 nuclease in mammalian cells with an expanded PAM compatibility.

### StaCas9 displays robust genome-editing activity across endogenous loci

We next evaluated the genome-editing performance of StaCas9 at a panel of 30 endogenous genomic loci bearing NNGN PAMs in HEK293T cells. SpCas9-NG, which is capable of targeting the same loci, was included as a reference nuclease. To ensure comparable expression levels, both nucleases were expressed from an identical vector backbone (Fig. 2A). Genomic DNA was harvested seven days after transfection of Cas9- and sgRNA-expression plasmids and subjected to targeted deep sequencing.

**Figure 2.**
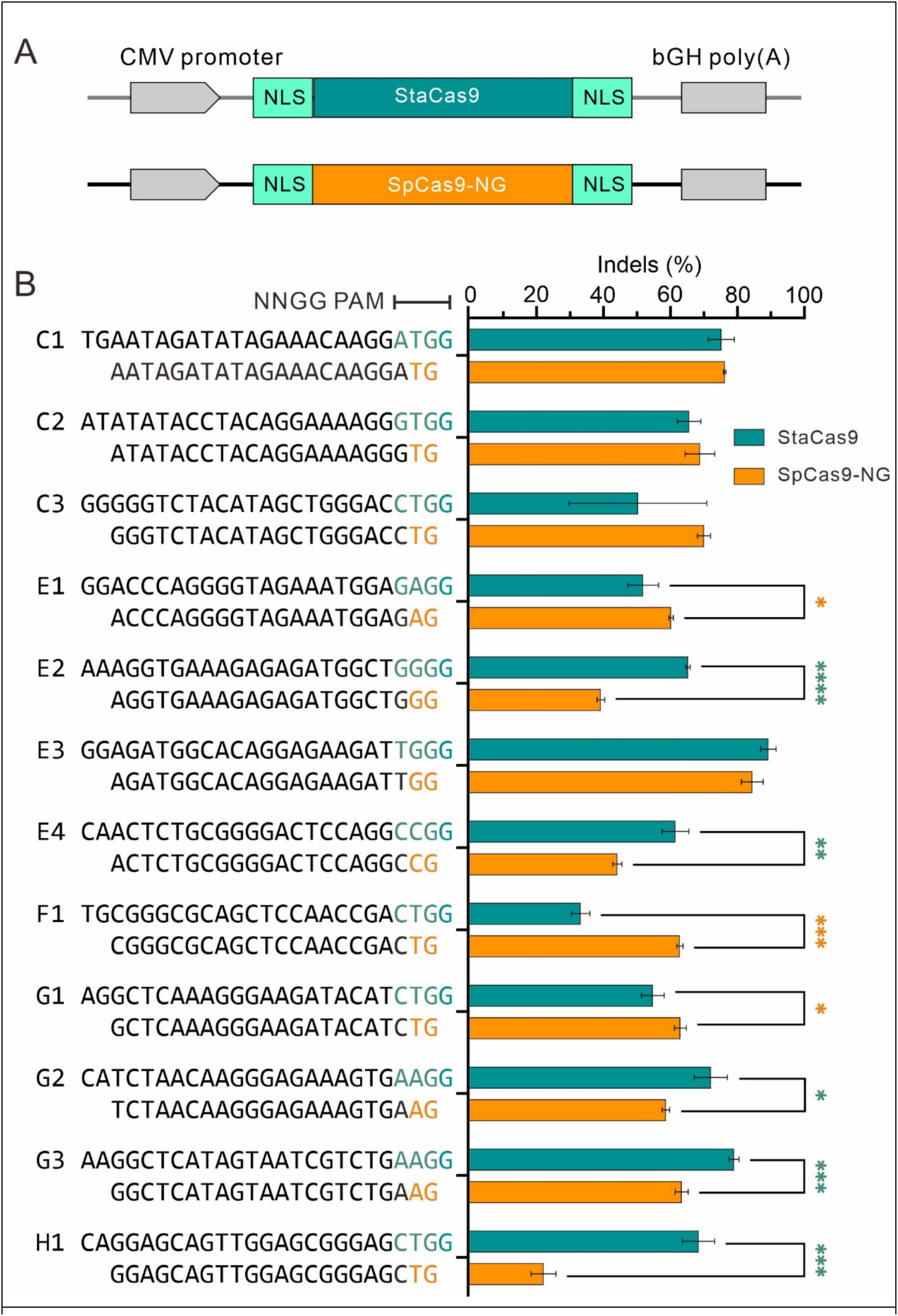
Evaluation of StaCas9 activity at endogenous targets with NNGG PAMs. **(A)** Schematic representation of the Cas9 expression plasmids used for comparative analysis. **(B)** Genome-editing efficiencies of StaCas9 and SpCas9-NG at a panel of 12 endogenous genomic loci containing NNGG PAMs in HEK293T cells. Target sequences are shown on the left. PAM sequences recognized by StaCas9 are highlighted in green, whereas PAM sequences recognized by SpCas9-NG are highlighted in orange. Data are presented as mean ± standard deviation (s.d.) from three independent biological replicates (n = 3). Statistical significance was assessed using a two-tailed statistical test with the following thresholds: *P < 0.05, **P < 0.01, ***P < 0.001, and ****P < 0.0001.

Both nucleases exhibited locus-dependent editing efficiencies. Among the 12 loci containing NNGG PAMs, StaCas9 achieved editing efficiencies of up to 95% (Fig. 2B) and showed comparable or higher activity than SpCas9-NG at nine loci. In contrast, at loci containing NNGA, NNGC, or NNGT PAMs, StaCas9 generally exhibited reduced editing efficiencies, outperforming or performing comparably to SpCas9-NG at only four of the 18 loci tested (Fig. S6A–C).

To further assess cell-type dependence, StaCas9 activity was examined in additional human cell lines (HCT116, SH-SY5Y, and A375) as well as mouse N2a cells. Editing efficiencies varied across both target loci and cell types; however, a consistent preference for NNGG PAMs was observed across all tested cell lines (Fig. S7A-B and S8A-B). A modest preference for NNGA PAMs was additionally detected in SH-SY5Y and N2a cells, whereas little activity was observed at NNGC and NNGT PAMs. Among 16 endogenous loci containing NNGC or NNGT PAMs, StaCas9 outperformed or performed comparably to SpCas9-NG at nine loci (Fig. S7A-B and S8A-B).

### Assessment of StaCas9 genome-editing specificity

To assess the mismatch tolerance of StaCas9, we employed a GFP-activation assay using a fixed CTGG PAM. A panel of ten sgRNAs carrying dinucleotide mismatches at distinct positions along the protospacer was designed, with the fully matched sgRNA serving as a control. Editing efficiency was quantified based on the percentage of GFP-positive cells. Editing analysis revealed that StaCas9 tolerated dinucleotide mismatches to a limited extent at positions 10 and 16, whereas mismatches at other positions resulted in a pronounced loss of activity, indicating stringent target recognition (Fig. 3A).

**Figure 3.**
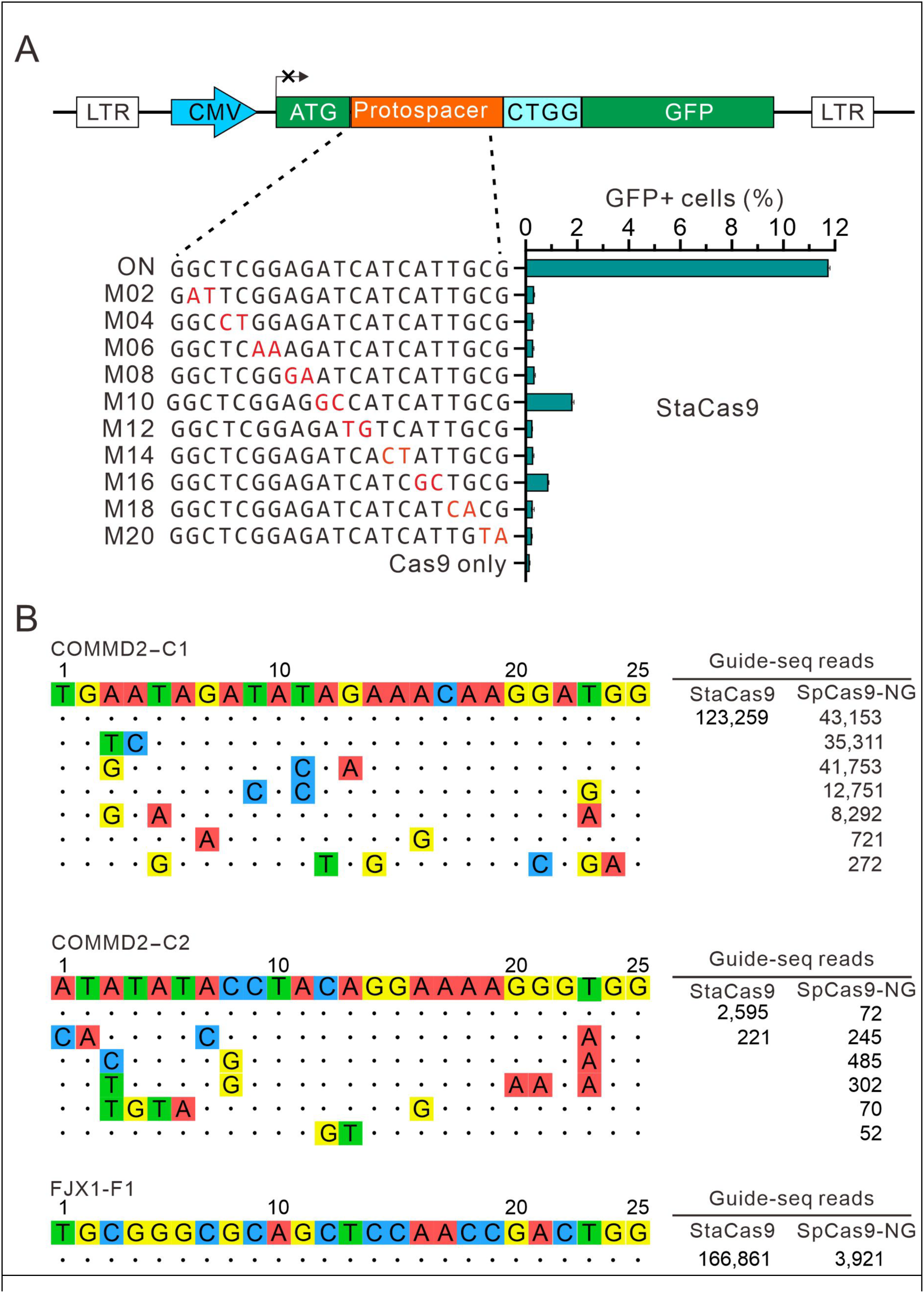
Specificity characterization of StaCas9. **(A)** Assessment of StaCas9 mismatch tolerance using a GFP-activation reporter assay. A schematic of the GFP reporter construct is shown at the top. Below, sgRNAs containing dinucleotide mismatches (highlighted in red) at different positions along the protospacer are depicted. Editing efficiency was quantified as the percentage of GFP-positive cells (n = 3 independent biological replicates). **(B)** Genome-wide off-target analysis of StaCas9 and SpCas9-NG using GUIDE-seq. On-target and detected off-target sequences are shown on the left, with GUIDE-seq read counts shown on the right. Mismatches relative to the on-target sequence are indicated in colour.

We next examined genome-wide off-target activity using GUIDE-seq ^19^ in HEK293T cells, with SpCas9-NG included as a comparator nuclease. Two sgRNAs targeting the COMMD2 locus and one sgRNA targeting the FJX1 locus were evaluated. Cas9 expression plasmids, sgRNA expression constructs, and GUIDE-seq oligonucleotides were co-electroporated into cells, followed by genomic DNA isolation five days later for sequencing-based analysis. GUIDE-seq profiling revealed strong on-target cleavage signals for both nucleases at the COMMD2-C1 site (Fig. 3B). Notably, six off-target sites were detected for SpCas9-NG, whereas no off-target sites were identified for StaCas9 at this locus. At the COMMD2-C2 site, StaCas9 generated a single detectable off-target event, compared with five off-target sites observed for SpCas9-NG. No off-target cleavage was detected for either nuclease at the FJX1-F1 site. Together, these results indicate that StaCas9 exhibits stringent sequence discrimination and high genome-wide specificity in mammalian cells.

### Therapeutic potential of StaCas9 for editing PCSK9

To assess the therapeutic potential of StaCas9, we evaluated its ability to edit the PCSK9 gene in HEK293T cells, with SpCas9-NG included as a comparator. A panel of 12 endogenous target sites that could be recognized by both nucleases was designed (Fig. 4A). Seven days after transfection with Cas9- and sgRNA-expression plasmids, genomic DNA was isolated and analyzed by targeted deep sequencing. Both nucleases achieved robust editing, with maximal efficiencies approaching 70% at certain loci (Fig. 4B). StaCas9 exhibited higher editing efficiencies than SpCas9-NG at three target sites, all of which contained NNGA or NNGG PAMs. In contrast, SpCas9-NG outperformed StaCas9 at seven sites, predominantly those harboring NNGC or NNGT PAMs. When editing efficiencies across all tested sites were considered collectively, SpCas9-NG showed a modestly higher overall activity; however, this difference did not reach statistical significance (Fig. 4C). These results highlight StaCas9 as a promising option for applications requiring high specificity and robust activity at NNG PAMs, particularly in size-limited delivery settings such as AAV.

**Figure 4.**
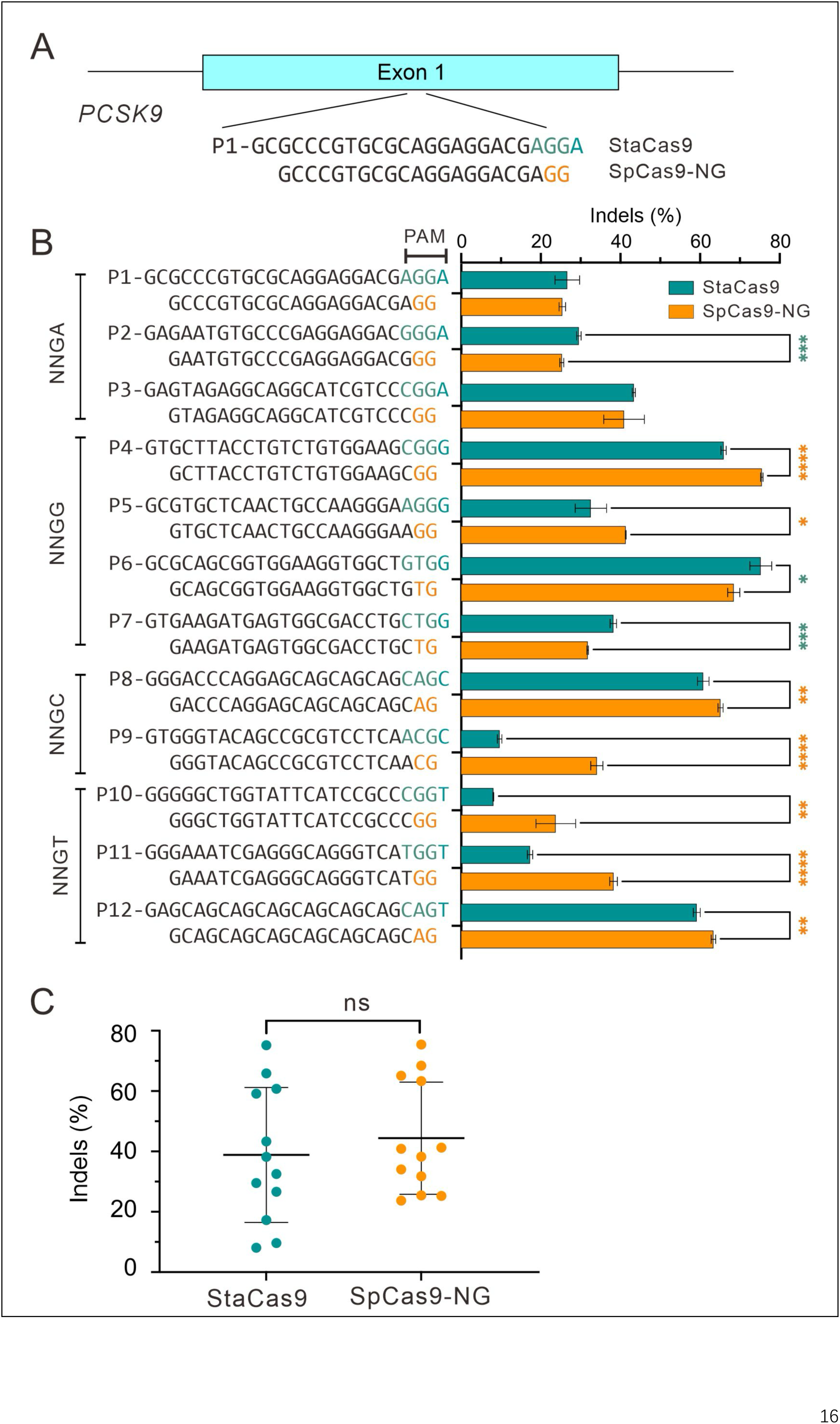
Therapeutic potential of StaCas9 for PCSK9 editing. **(A)** Schematic of sgRNA design targeting PCSK9. The structure of PCSK9 exon 1 is shown at the top, with representative target sites that can be recognized by both StaCas9 and SpCas9-NG indicated below. **(B)** Genome-editing activities of StaCas9 and SpCas9-NG at a panel of 12 endogenous PCSK9 target sites in HEK293T cells. Data are presented as mean ± standard deviation (s.d.) from three independent experiments (n = 3). **(C)** Quantification of indel frequencies for both nucleases, determined by targeted deep sequencing. Statistical significance was assessed using a two-tailed statistical test with the following thresholds: *P < 0.05, **P < 0.01, ***P < 0.001, and ****P < 0.0001

### Engineering StaCas9 to relax nucleotide preference at PAM position 4

Sequence logo and PAM wheel analyses indicated that StaCas9 exhibits a mild preference for A or G at the fourth position of the PAM (Fig. 1D–E). Consistent with this observation, StaCas9 generally showed reduced editing efficiency at targets bearing NNGC or NNGT PAMs (Fig. S5A–C).

Previously, we evolved a SlugCas9 variant with relaxed NNG PAM recognition (SlugCas9-NNG) using phage-assisted continuous evolution ^20^. This variant harbors six amino acid substitutions (Q782R, S888R, L906R, N984S, E1012K, and K1016I) that collectively broaden its PAM compatibility. To transfer this strategy to StaCas9, we aligned the amino acid sequences of StaCas9 and SlugCas9-NNG and identified the corresponding residues in StaCas9 (Q781, E887, L905, S983, S1010, and K1014; Fig. S9A). Because StaCas9 already contains a serine at position 983, corresponding to the N984S substitution in SlugCas9-NNG, no modification was introduced at this site. The remaining five residues were mutated to match those in SlugCas9-NNG, generating the engineered variant StaCas9-NG.

Structural modeling suggested that these substitutions enable additional hydrogen-bond interactions with the PAM duplex, potentially accounting for the relaxed PAM recognition (Fig. S9B–C). We next characterized the PAM profile of StaCas9-NG using the GFP-activation reporter assay (Fig. S10A). Co-transfection of StaCas9-NG and its corresponding sgRNA into reporter cells led to GFP activation (Fig. S10B). Deep sequencing of the randomized PAM region revealed that StaCas9-NG recognizes an NNG PAM with reduced nucleotide bias at position 4 (Fig. S10C–D).

To assess genome-editing activity, we tested StaCas9-NG at a panel of 16 endogenous loci, with wild-type StaCas9 serving as a reference. StaCas9-NG achieved editing efficiencies of up to ∼30%, whereas wild-type StaCas9 reached efficiencies as high as ∼90% at the same targets (Fig. S11A). Overall, StaCas9-NG exhibited significantly reduced editing activity compared with the wild-type enzyme (Fig. S11B). Together, these results demonstrate that StaCas9-NG achieves relaxed PAM recognition at the expense of genome-editing efficiency.

### Engineering StaCas9 to enhance activity

To enhance the editing activity of StaCas9, we employed an automated computational pipeline for rational protein design and generated a panel of 45 candidate point mutations. Each mutation was individually introduced into StaCas9 and evaluated using a GFP-independent genome-editing assay targeting the EMX1 locus in HEK293T cells. Notably, five variants, E887R, S998K, E420R, G925R, and I817R, exhibited significantly improved activity, achieving up to a 1.63-fold increase compared with wild-type StaCas9 (Fig. 5A).

**Figure 5.**
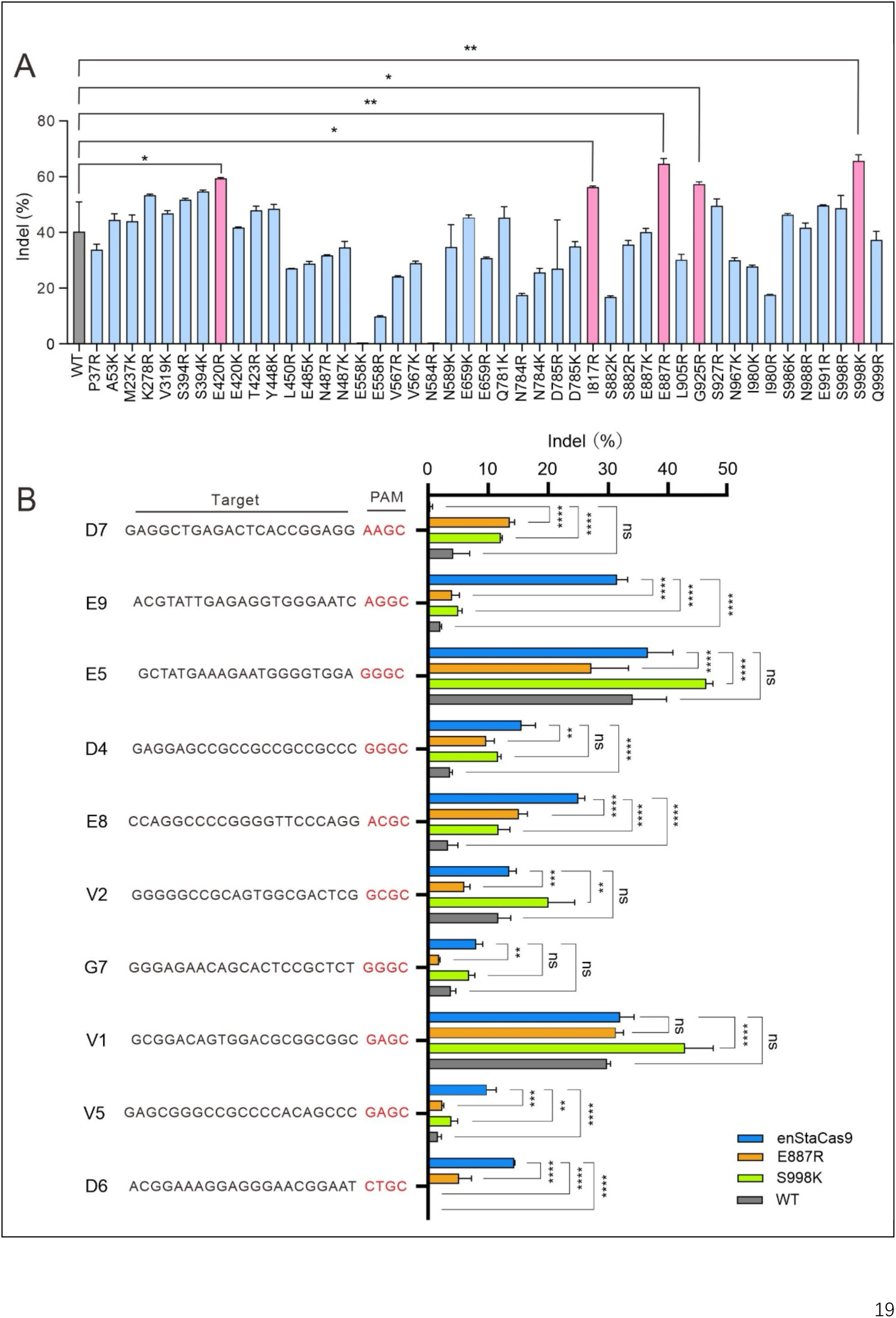
Engineering StaCas9 to enhance genome-editing activity. **(A)** Screening of rationally designed StaCas9 variants for enhanced activity. Forty-five single–amino acid substitutions were individually introduced into StaCas9 and evaluated using an endogenous EMX1 target site in HEK293T cells. Genome-editing efficiencies were quantified by targeted deep sequencing. Five mutations (E887R, S998K, E420R, G925R, and I817R) significantly increased editing activity relative to wild-type StaCas9. **(B)** Evaluation of combinatorial mutations. The two most effective substitutions, E887R and S998K, were combined to generate an enhanced variant termed enStaCas9. Genome-editing efficiencies of wild-type StaCas9, single mutants (E887R or S998K), and enStaCas9 were assessed across six endogenous target sites in HEK293T cells. enStaCas9 exhibited higher activity than wild-type StaCas9 at four of ten loci. Data are presented as mean ± s.d. from three independent biological replicates (n = 3). Statistical significance was determined using a two-tailed test (*P < 0.05, **P < 0.01, ***P < 0.001, ****P < 0.0001).

To evaluate whether the beneficial mutations act additively, the two top-performing substitutions, E887R and S998K, were combined to generate an enhanced variant, designated enStaCas9. We then assessed the activity of enStaCas9 across ten endogenous genomic targets. Compared with wild-type StaCas9, enStaCas9 exhibited higher editing efficiency at five loci; relative to the single E887R mutant, enStaCas9 showed improved activity at eight loci; and compared with the single S998K mutant, enStaCas9 outperformed four loci (Fig. 5B). These results demonstrate that rational, structure-guided engineering can synergistically enhance StaCas9 activity in mammalian cells.

Structural modeling was used to explore the potential mechanisms underlying the enhanced activity of the E887R and S998K mutations. Residue E887 is positioned adjacent to the DNA backbone within the protospacer region (Fig. S12). Replacement of glutamate with arginine introduces a positively charged side chain that is oriented toward the DNA phosphate backbone, with a predicted distance of ∼3.6 Å, consistent with the formation of stabilizing electrostatic interactions.

Residue S998 is located near the PAM-proximal DNA region. Substitution with lysine introduces a longer, positively charged side chain that is also directed toward the DNA backbone, potentially reinforcing local protein–DNA contacts (Fig. S12). Together, these structural features suggest that the E887R and S998K mutations may enhance Cas9–DNA interactions, providing a structural basis for the increased genome-editing activity observed for enStaCas9.

## Discussion

The diversity of PAM recognition among Cas9 nucleases is a central determinant of their genome-editing versatility. Mining naturally occurring Cas9 enzymes with distinct PAM specificities has proven to be an efficient strategy for expanding the editable genome. Using this approach, we and others have developed multiple type II-A Cas9 nucleases, including ScCas9 ^21^, SauriCas9 ^14^, SeqCas9 ^22^, and Tsp2Cas9 ^23^, as well as several type II-C Cas9 nucleases such as GeoCas9 ^24^, Nme2Cas9 ^25^, and Nsp2Cas9 ^26^. Beyond Cas9, diverse Cas12 nucleases with distinct PAM requirements, such as AsCas12a ^27^, AaCas12b ^28^, Plm2Cas12e ^29^, AsCas12f ^30, 31^, Un1Cas12f1 ^32^, and Cas12j-8 ^33^, have also been harnessed for genome editing.

In this study, we identify StaCas9 as a naturally evolved type II-A Cas9 nuclease with a distinct preference for NNG PAMs, thereby further expanding the functional space of compact Cas9 enzymes suitable for therapeutic applications. Using an unbiased GFP-based PAM screening assay, we demonstrate that StaCas9 recognizes a relaxed NNG PAM, with pronounced enrichment for NNGG and NNGA motifs, in sharp contrast to the stringent NNGRRT PAM requirement of SaCas9 ^8^. Structural and sequence analyses further identify S983 within the PAM-interacting (PI) domain as a key determinant of this altered specificity. Unlike the conserved asparagine at the corresponding position in SaCas9, S983 in StaCas9 is predicted to remodel local interactions within the PAM duplex, providing a mechanistic link between natural sequence variation and functional diversification of PAM recognition. Notably, our previous study identified three residues that govern the NNGG PAM requirement of SaCas9, consistent with the present findings and supporting the notion that variation at a limited number of key positions can fine-tune PAM preference across SaCas9 orthologs ^34^. Together, these findings underscore how subtle changes within the PI domain can drive meaningful diversification of PAM preference and reinforce the value of mining natural Cas9 diversity.

Beyond PAM recognition, our results reveal that genome-editing activity is strongly influenced by sequence context, even among Cas9 nucleases targeting the same PAM class. StaCas9 exhibits robust editing activity at a subset of endogenous loci, particularly those bearing NNGG PAMs, where it matches or surpasses the performance of SpCas9-NG, while showing reduced activity at NNGC and NNGT sites. This sequence-dependent activity profile is consistently observed across multiple cell types, indicating that it reflects intrinsic biochemical properties rather than cell-specific effects. Similar sequence-dependent behavior is also evident for other Cas9 nucleases, such as SpCas9 ^35^, SpeCas9 ^22^, and Nme2Cas9 ^25^.

These findings highlight a key principle for therapeutic genome editing: no single Cas9 nuclease is universally optimal, even when PAM compatibility overlaps. Instead, each Cas9 enzyme occupies a distinct activity landscape shaped by PAM sub-preference, protospacer sequence, and local genomic context. In this regard, StaCas9 should be viewed not as a replacement for previously developed NNG-recognizing Cas9 nucleases, but as a complementary genome-editing tool that excels at specific target classes.

## Materials and methods

### Cas9 orthologs expression plasmid construction

The plasmid pAAV-CMV-puro backbone was obtained by digesting plasmid pAAV-CMV-SauriCas9-puro (Addgene No.135965) with restriction enzymes AgeI-HF (NEB, USA) and BamHI-HF (NEB, USA). The human codon-optimized Cas9 gene was synthesized by GENEWIZ (Suzhou, China) and cloned into the pAAV-CMV-puro backbone by the Instant Sticky-end Ligase (NEB, USA). The sequence of each Cas9 protein was confirmed by Sanger sequencing (GENEWIZ, China). The sequences of the Cas9 gene are listed in Table S1.

The sgRNA was ligated into the backbone by digesting plasmid pAAV-CMV-StaCas9-puro with restriction enzyme BsaI-HF (NEB, USA) to obtain the sgRNA-expressing plasmid. The primer sequences and target sequences are listed in Table S2 and Table S3, respectively.

The sgRNA fragments were synthesized by GENEWIZ (Suzhou, China) and the vector were PCR-amplified using KOD DNA polymerase and subsequently assembled via the NEBuilder HiFi DNA Assembly Cloning Kit to obtain the sgRNA-expressing plasmid. The sgRNA sequences and primers are listed in Table S1 and Table S2, respectively.

The plasmid pAAV-CMV-StaCas9-puro was linearized by inverse PCR using KOD DNA polymerase (TOYOBO), with the amino acid mutations designed at the 5′ ends of the primers. These mutations were subsequently introduced into the plasmid using the NEBuilder HiFi DNA Assembly Cloning Kit (NEB, USA) to generate the StaCas9 mutant plasmid. The same approach was employed to construct the pAAV-CMV-StaCas9-puro mutant plasmid. Table S2 and Table S3 provide the sgRNA sequences and primer sequences, respectively.

### Cell culture and transfection

HEK 293T, A375, SH-SY5Y, and N2a cells were cultured in Dulbecco’s Modified Eagle’s Medium (DMEM, Gibco, USA). HCT116 cells were maintained in McCoy’s 5A. All the cells were supplemented with 10% FBS (Gibco, USA), 100 U/mL penicillin, and 100 mg/mL streptomycin (Gibco, USA) in humidified, 37 °C chamber with 5% CO_2_. All cell lines used tested negative for Mycoplasma contamination.

HEK293T, and N2a cells were transfected with Lipofectamine 2000 (Life Technologies, USA) according to the manufacturer’s instructions. SH-SY5Y, A375, and HCT116 cells were transfected with Lipofectamine 3000 (Life Technologies, USA) according to the manufacturer’s instructions. For Cas9 PAM sequence screening, HEK293T cells were plated into 10 cm dishes and transfected at 50–60% confluency with a total of 10 μg of Cas9 plasmid and 5 μg of sgRNA plasmid in 10 cm dishes. For genome editing of Cas9, HEK293T, N2a, SH-SY5Y, A375, and HCT116 cells were seeded into a 24-well plate and transfected with Cas9-sgRNA plasmids (1 μg) by Lipofectamine (2 μL) Cells were collected five days after transfection. Genomic DNA was isolated, and the target sites were PCR-amplified by nested PCR amplification and purified by a Gel Extraction Kit (QIAGEN, USA) for deep sequencing. The primer sequences are listed in Table S3.

### PAM Sequence Analysis

Twenty-base-pair sequences (AAGCCTTGTTTGCCACCATG/GTGAGCAAGG GCGAGGAGCT) flanking the target sequence (GAACGGCTCGGAGATCATC ATTGCGNNNNNNN) were used to fix the target sequences. GCG and GTGAGCAAGGGCG AGGAGCT were used to fix a 7-bp random sequence. Target sequences with in-frame mutations were used for PAM analysis. The 7-bp random sequence was extracted and visualized by WebLogo and a PAM wheel chart to identify PAMs.

### Flow cytometry analysis

Transfected library cells with a certain percentage of GFP-positive cells were collected by centrifugation at 1000 rpm for 5 min and resuspended in PBS. Then, GFP-positive cells were collected by flow cytometry (SONY, Japan) and cultured in six-well plates. Five days after culture, the genomic DNA was isolated for deep sequencing.

### Test of Cas9 specificity

To test the specificity of StaCas9, we generated a GFP-reporter cell line with 5’-GTGG PAM. HEK293T cells were seeded into 24-well plates and transfected with StaCas9-sgRNA plasmids (1 μg) by using Lipofectamine 2000 (2 μL). Five days after editing, the GFP-positive cells were analyzed on the Calibur instrument (Cytek, USA). Data were analyzed using the FlowJo software.

### GUIDE-Seq

GUIDE-seq experiments were performed as described previously, with minor modifications. Briefly, 2×10^5^ HEK293T cells were transfected with 1000 ng of StaCas9-sgRNA plasmids, and 100 pmol of annealed GUIDE-seq oligonucleotides by electroporation and then seeded into 6 wells. The electroporation voltage, width, and the number of pulses were 1,500 V, 30 ms, and 1 pulse, respectively. Genomic DNA was extracted with the DNeasy Blood and Tissue kit (QIAGEN, USA) 6 days after transfection according to the manufacturer’s protocol. The genome library was prepared and subjected to deep sequencing.

### Automated computational pipeline for rational protein design

To enhance the editing activity of StaCas9, we implemented an automated computational pipeline for structure-guided protein engineering ^36^. First, protein–DNA–RNA ternary complex structures were modeled based on multiple target sites with varying editing efficiencies using AlphaFold3 ^37^. For each complex, PyMOL (http://www.pymol.org/pymol) was used to measure spatial distances between amino acid side chains and the DNA backbone, and residues with close proximity (< 6.5 Å) were selected as candidate mutation sites.

Subsequent analyses focused on evaluating the electrostatic and structural impacts of mutations. Using the PyRosetta toolkit ^38^, we assessed changes in ionic and hydrogen bonding interactions upon mutation to arginine (R) or lysine (K), as only these positively charged residues were found to create new salt bridges with the DNA phosphate backbone. Mutations that introduced new favorable ionic interactions absent in the wild type were retained. Protein stability was evaluated with FoldX (v5.0) ^39^ by calculating the difference in folding free energy (ΔΔG) between wild-type and mutant structures. Mutations with ΔΔG > +3 kcal·mol⁻¹ were discarded to avoid destabilizing the protein fold. Solvent accessibility was further analyzed using PyRosetta to compute the solvent-accessible surface area (SASA) for each mutant. Mutations that increased SASA (ΔSASA > 0 Å²) were considered potentially beneficial for DNA interaction. Candidate mutations identified from four independent editing sites were integrated, and only residues that appeared in at least three of the four site-specific lists were selected for experimental validation.

### Quantification and statistical analysis

All the data are shown as mean ± SD. Statistical analyses were conducted using GraphPad Prism 9. Student’s *t*-test or one-way analysis of variance (ANOVA) was used to determine statistical significance between two or more groups, respectively. A value of *p* < 0.05 was considered to be statistically significant (\**p* < 0.05, \*\**p* < 0.01, \*\*\**p* < 0.001 , \*\*\*\**p* < 0.0001). All data needed to evaluate the conclusions in the paper are present in the paper and the Supplementary Materials.

## Author contributions

M.L., W.Y., Y.T., and J.L. performed experiments. M.L. and S.W. analyzed the data. Y.W. wrote the manuscript. Y.W. and J.L. provided experimental guidance.

## Acknowledgments

This work was supported by grants from the National Natural Science Foundation of China (82370254); the National Key Research and Development Program of China (2021YFA0910602); and the Science and Technology Research Program of Shanghai (24HC2810100, 23ZR1426000).

## Competing interests

Y.W. and J.L. have submitted patent applications on the basis of the results reported in this study. The remaining authors declare no competing interests.

