## Supplementary material for "Rational and computation-assisted engineering of a compact and efficient StaCas9 genome editor": StaCas9-sup.pdf

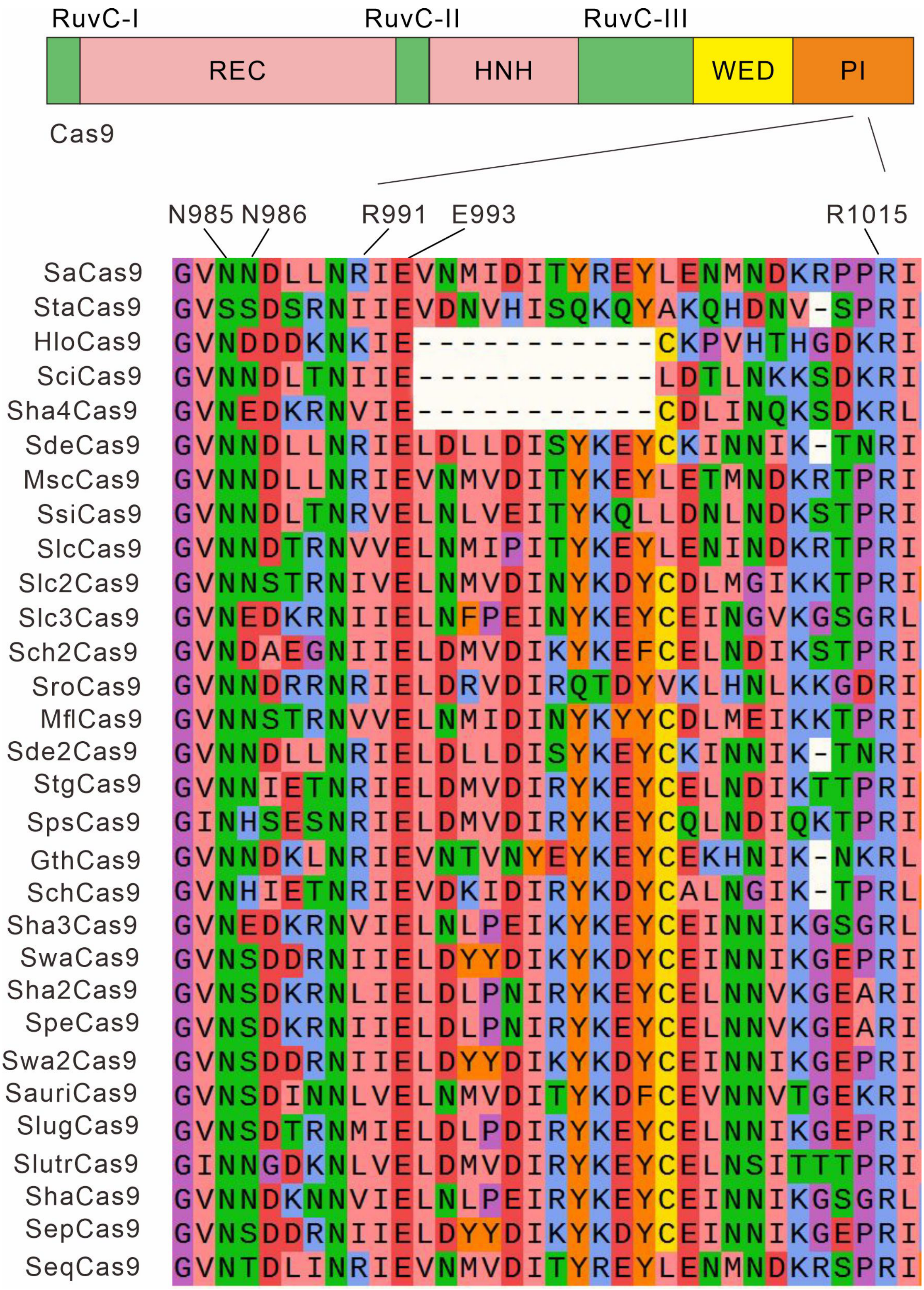


**Figure S1. Amino acid sequence alignment of SaCas9 orthologs in the PAM-interacting (PI) domain.** A schematic representation of Cas9 with annotated functional domains is shown at the top. The amino acid sequence alignment of the PI domain from 30 SaCas9 orthologs is shown below. Five residues known to play key roles in PAM recognition are highlighted above the alignment.

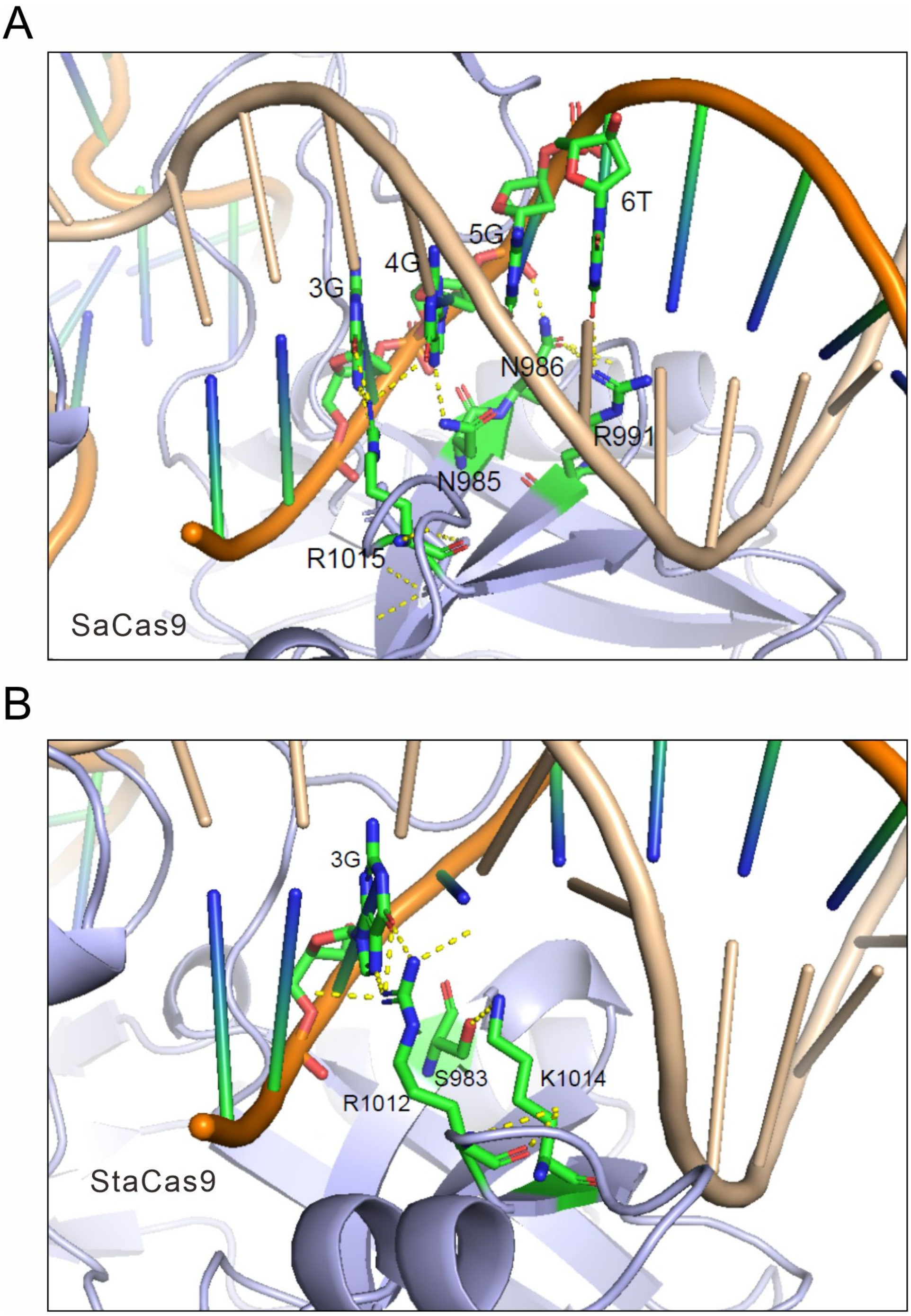


**Figure S2. AlphaFold3-predicted structural models of the PAM-interacting (PI) domains of SaCas9 and StaCas9.** (A) Predicted structure of the SaCas9 PI domain. Four residues involved in hydrogen-bond interactions with the PAM are highlighted. (B) Predicted structure of the StaCas9 PI domain. R1012 forms a hydrogen bond with the guanine at PAM position 3, whereas S983 forms a hydrogen bond with K1014 rather than directly contacting the PAM.

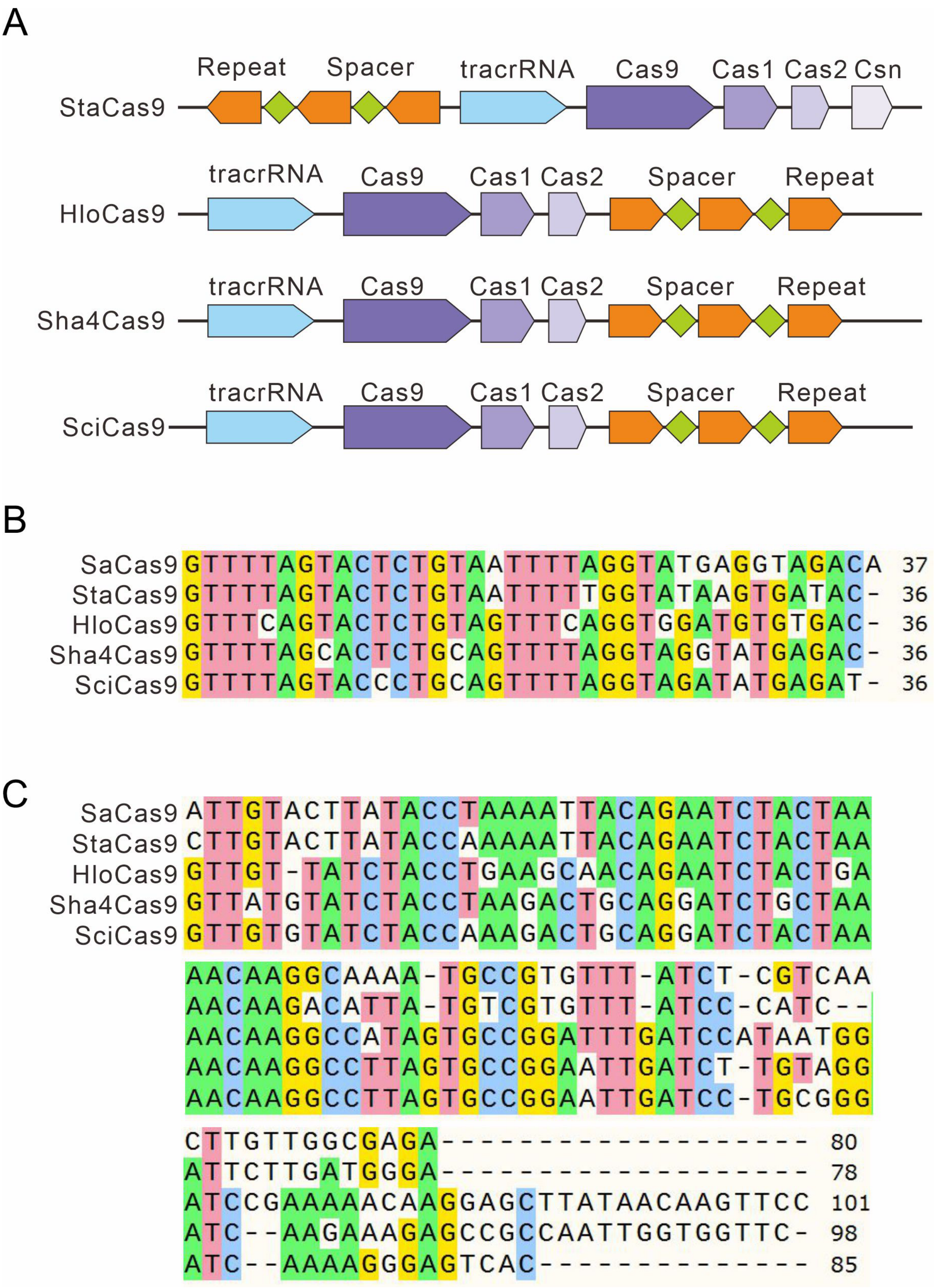


**Figure S3. Genomic organization and RNA sequence analysis of SaCas9 orthologs.** (A) Schematic representation of the genomic loci encoding the four SaCas9 orthologs. (B) Multiple sequence alignment of CRISPR repeat sequences from five SaCas9 orthologs. (C) Multiple sequence alignment of tracrRNA sequences from five SaCas9 orthologs.

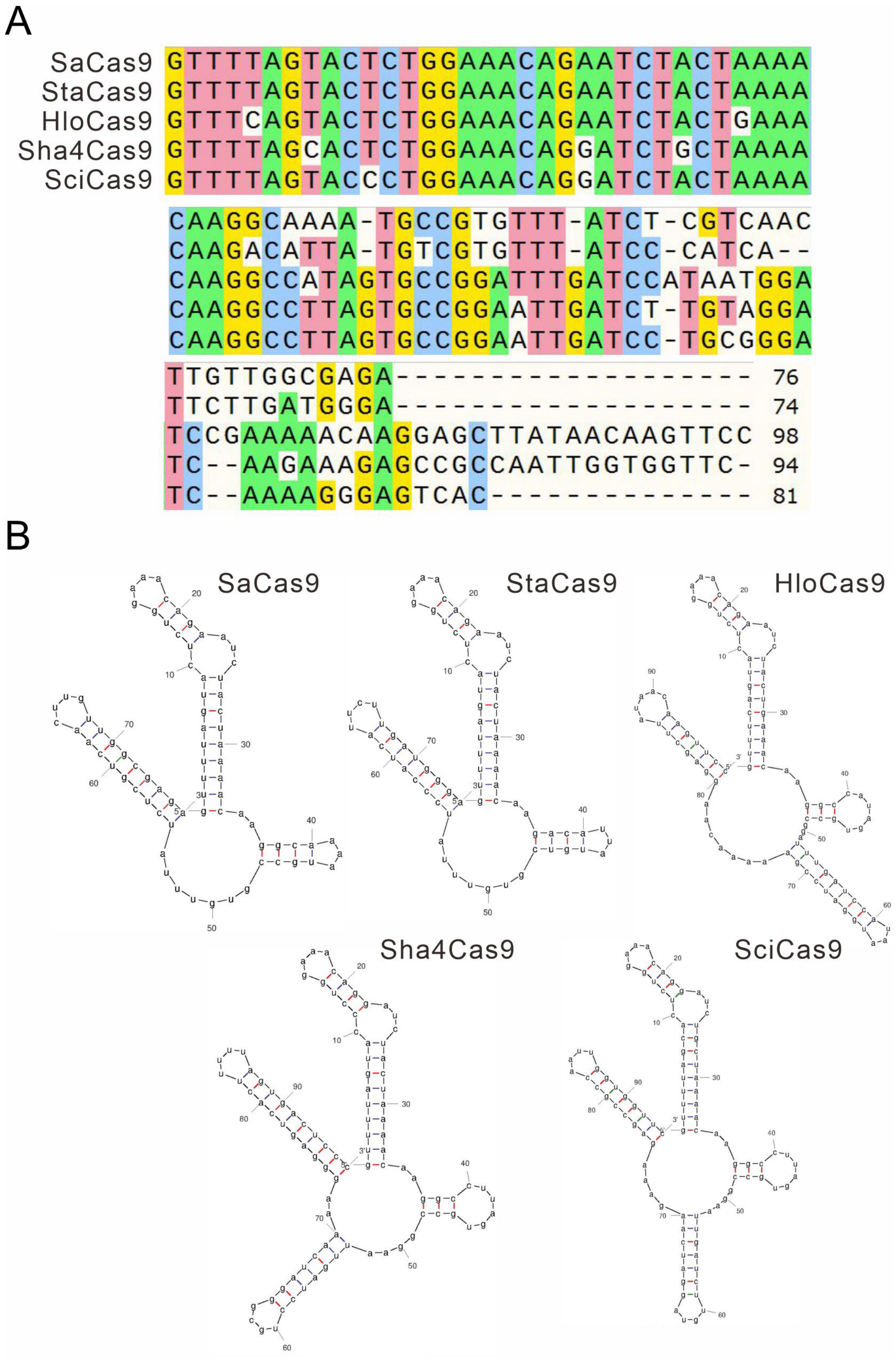


**Figure S4. sgRNA design and structural analysis of SaCas9 orthologs.** (A) Multiple sequence alignment of sgRNA scaffolds from five SaCas9 orthologs. (B) Predicted secondary structures of sgRNA scaffolds for SaCas9 orthologs, generated using the RNA Folding Form V2.3 online platform.

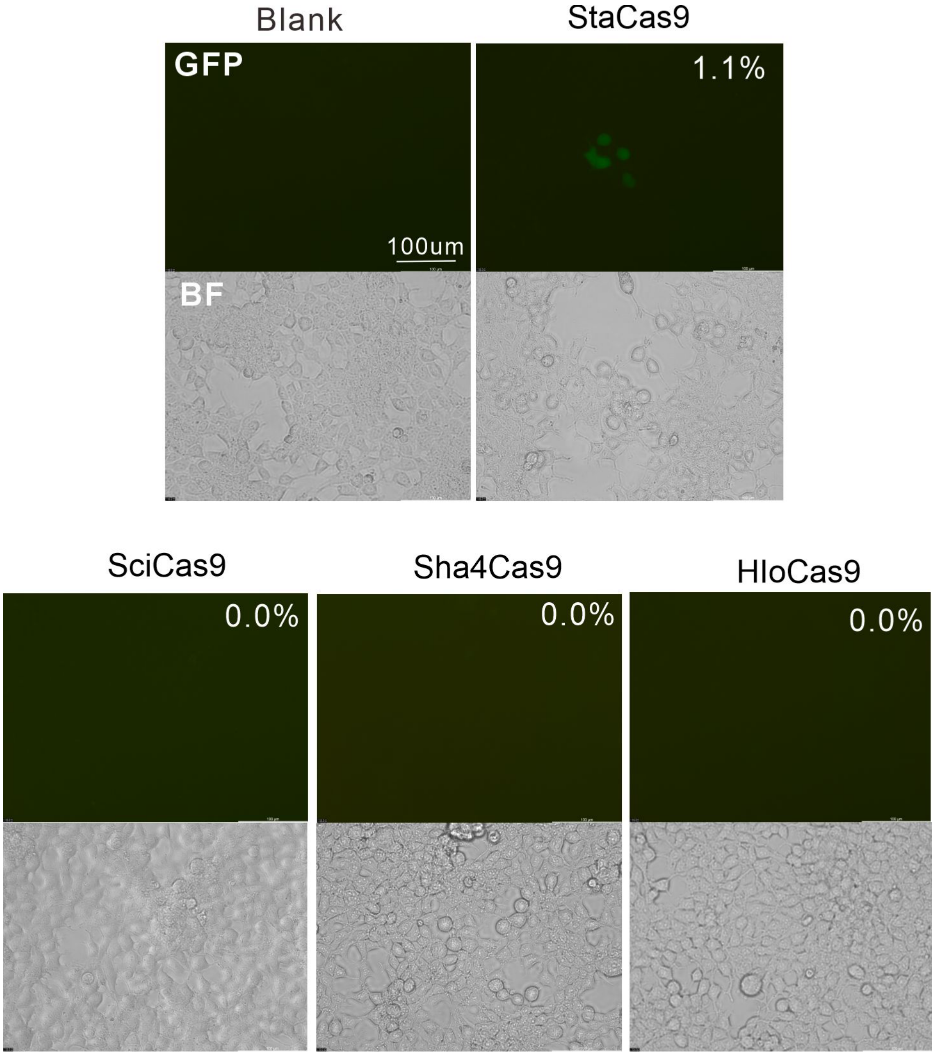


**Figure S5. Representative fluorescence images showing GFP activation induced by StaCas9 with its sgRNA scaffold.** SciCas9, Sha4Cas9, and HloCas9 did not induce GFP expression with their own sgRNA scaffold.

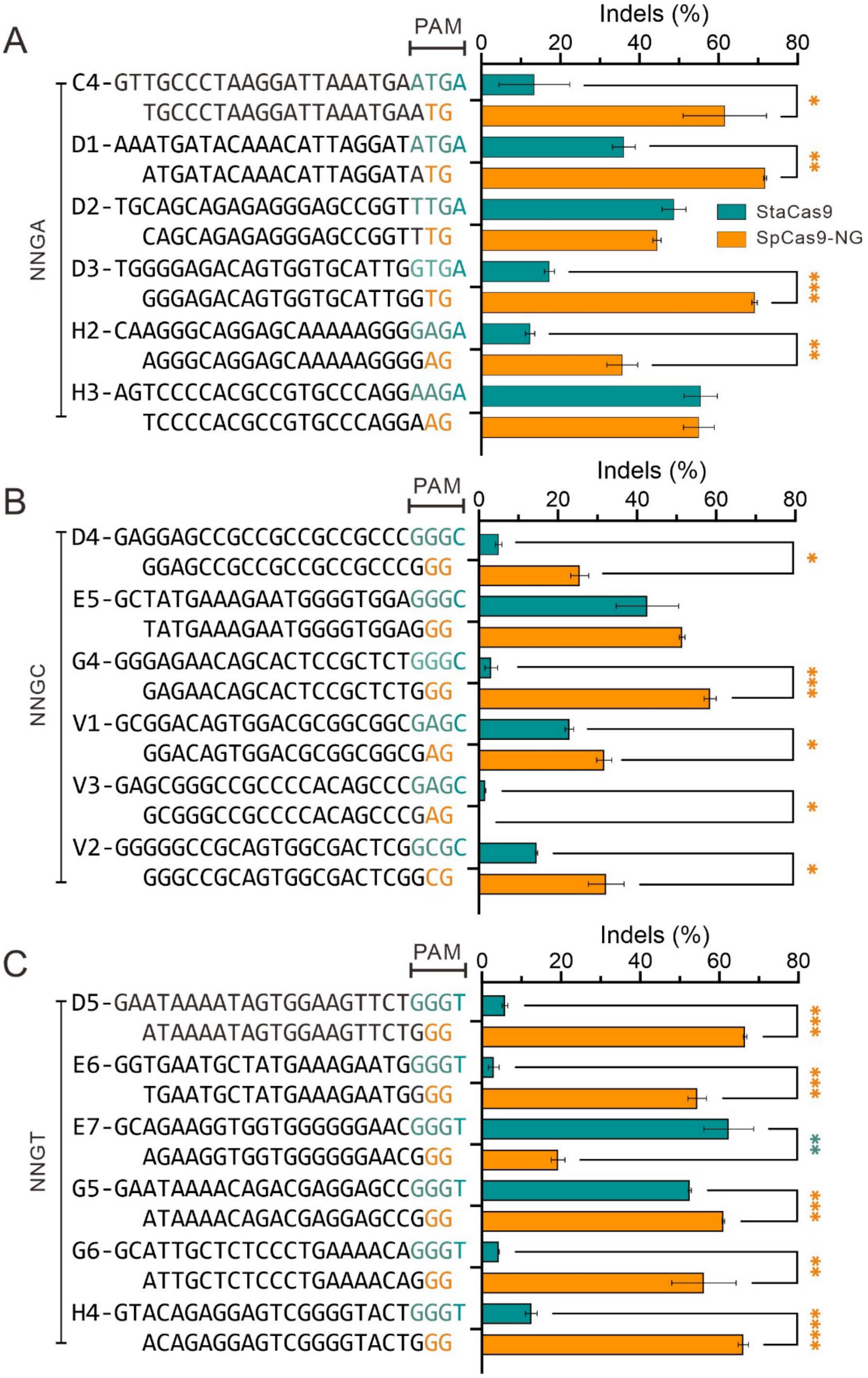


**Figure S6. Genome editing with StaCas9 and SpCas9-NG in a panel of 18 endogenous targets containing NNGA (A), NNGC (B), and NNGT (C) PAMs in HEK293T cells.** Data are presented as mean ± standard deviation (s.d.) from three independent biological replicates (n = 3). Statistical significance was assessed using a two-tailed statistical test with the following thresholds: *P < 0.05, **P < 0.01, ***P < 0.001, and ****P < 0.0001.

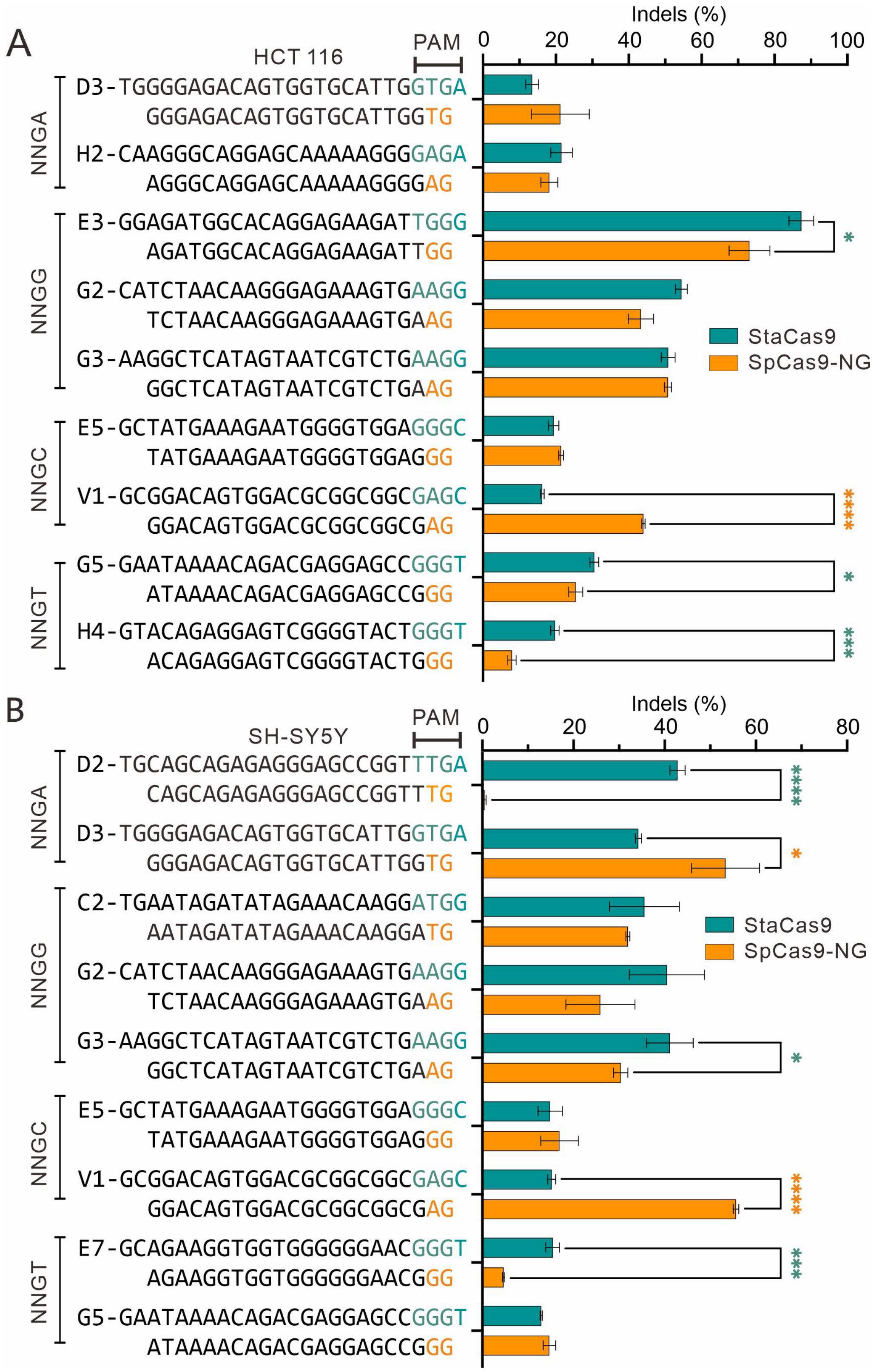


**Figure S7. StaCas9 enables genome editing in HCT116 (A) and SH-SY5Y (B) cells.** Data are presented as mean ± standard deviation (s.d.) from three independent biological replicates (n = 3). Statistical significance was assessed using a two-tailed statistical test with the following thresholds: *P < 0.05, **P < 0.01, ***P < 0.001, and ****P < 0.0001.

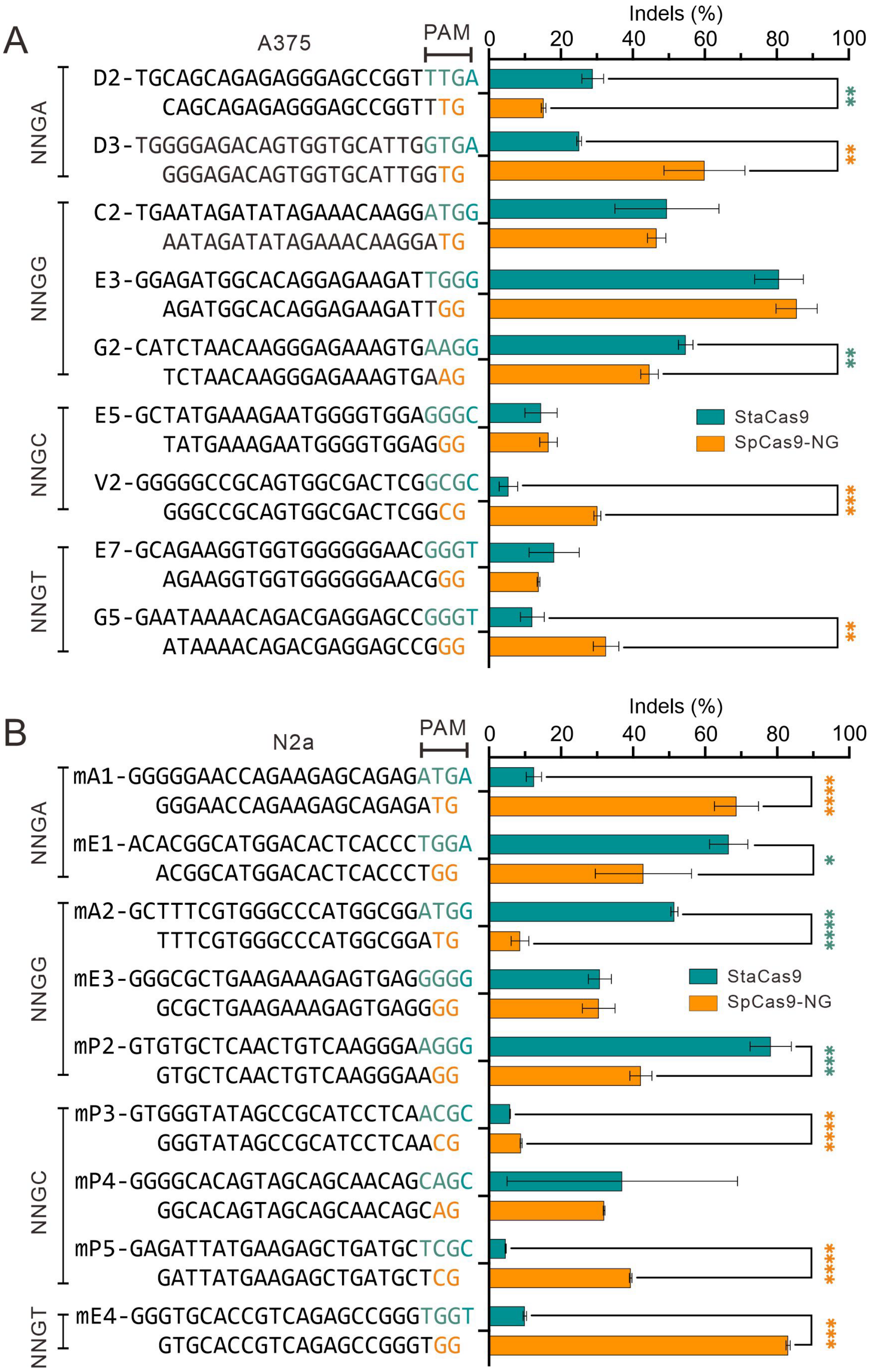


**Figure S8. StaCas9 enables genome editing in A375 (A) and N2a (B) cells.** Data are presented as mean ± standard deviation (s.d.) from three independent biological replicates (n = 3). Statistical significance was assessed using a two-tailed statistical test with the following thresholds: *P < 0.05, **P < 0.01, ***P < 0.001, and ****P < 0.0001.

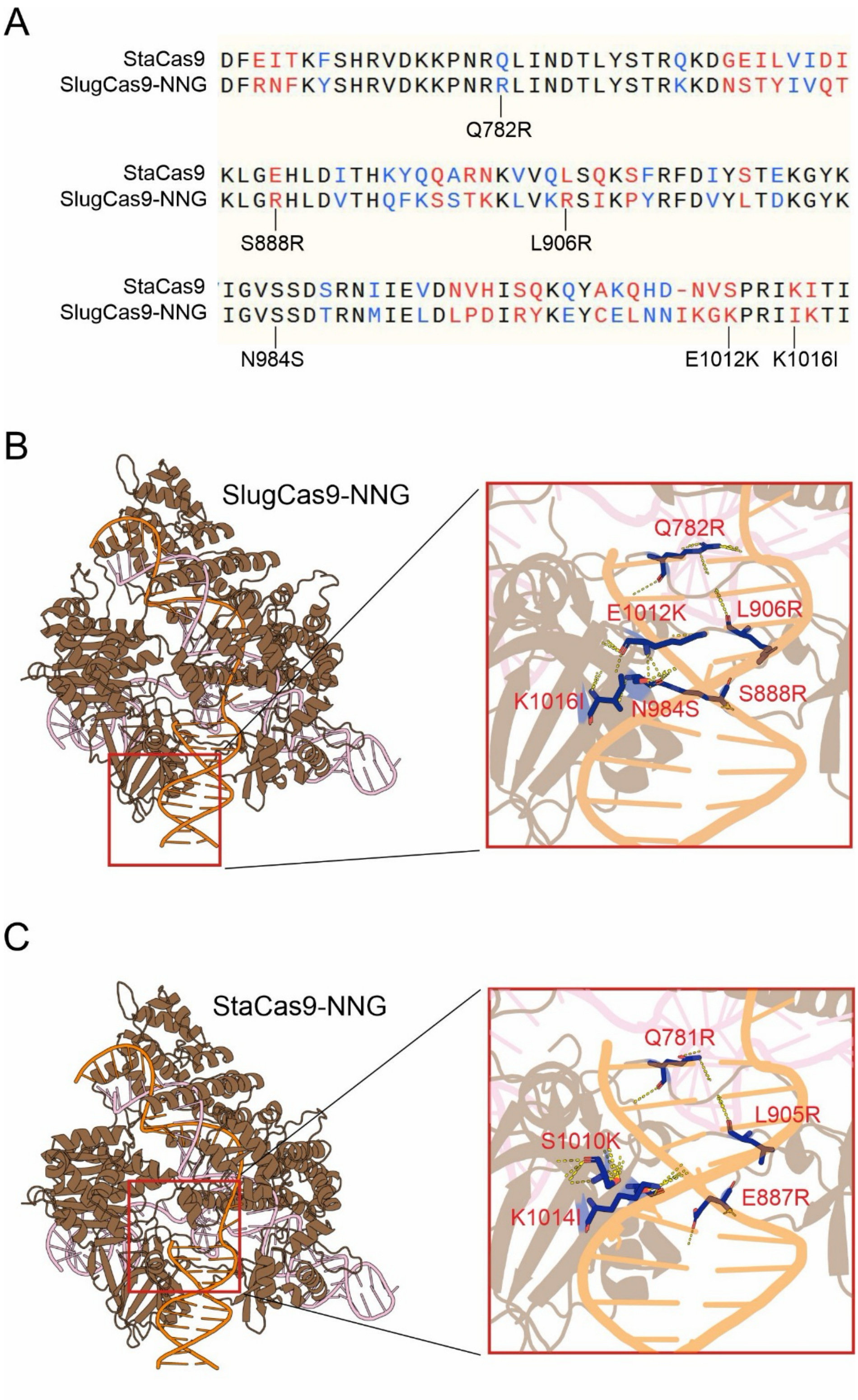


**Figure S9. Engineering of StaCas9-NNG for relaxed PAM recognition.** (A) Amino acid sequence alignment of StaCas9 and SlugCas9-NNG. Residues previously shown to be critical for relaxed PAM recognition in SlugCas9-NNG are indicated. (B, C) AlphaFold3-predicted structural models of SlugCas9-NNG (B) and StaCas9-NNG (C). Residues corresponding to the PAM-relaxing mutations are highlighted in red.

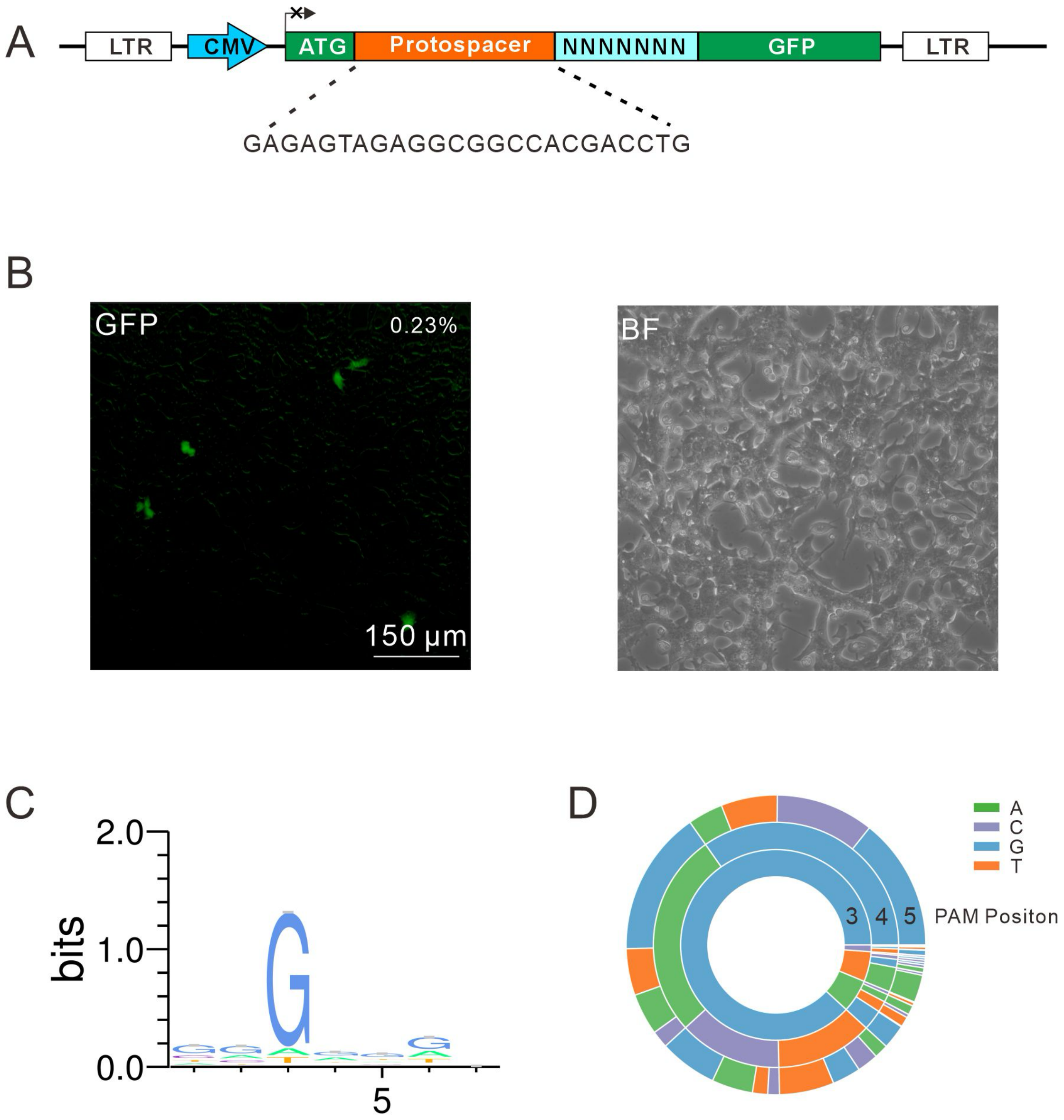


**Figure S10. PAM characterization of StaCas9-NNG using a GFP-activation assay.** (A) Schematic of the GFP-activation reporter construct. A target sequence (protospacer) followed by an 8-bp randomized sequence is inserted into the GFP coding region. (B) StaCas9-NNG induces GFP expression in reporter cells. (C) WebLogo representations derived from deep sequencing of edited target sites. (D) PAM wheel analysis based on deep sequencing data.

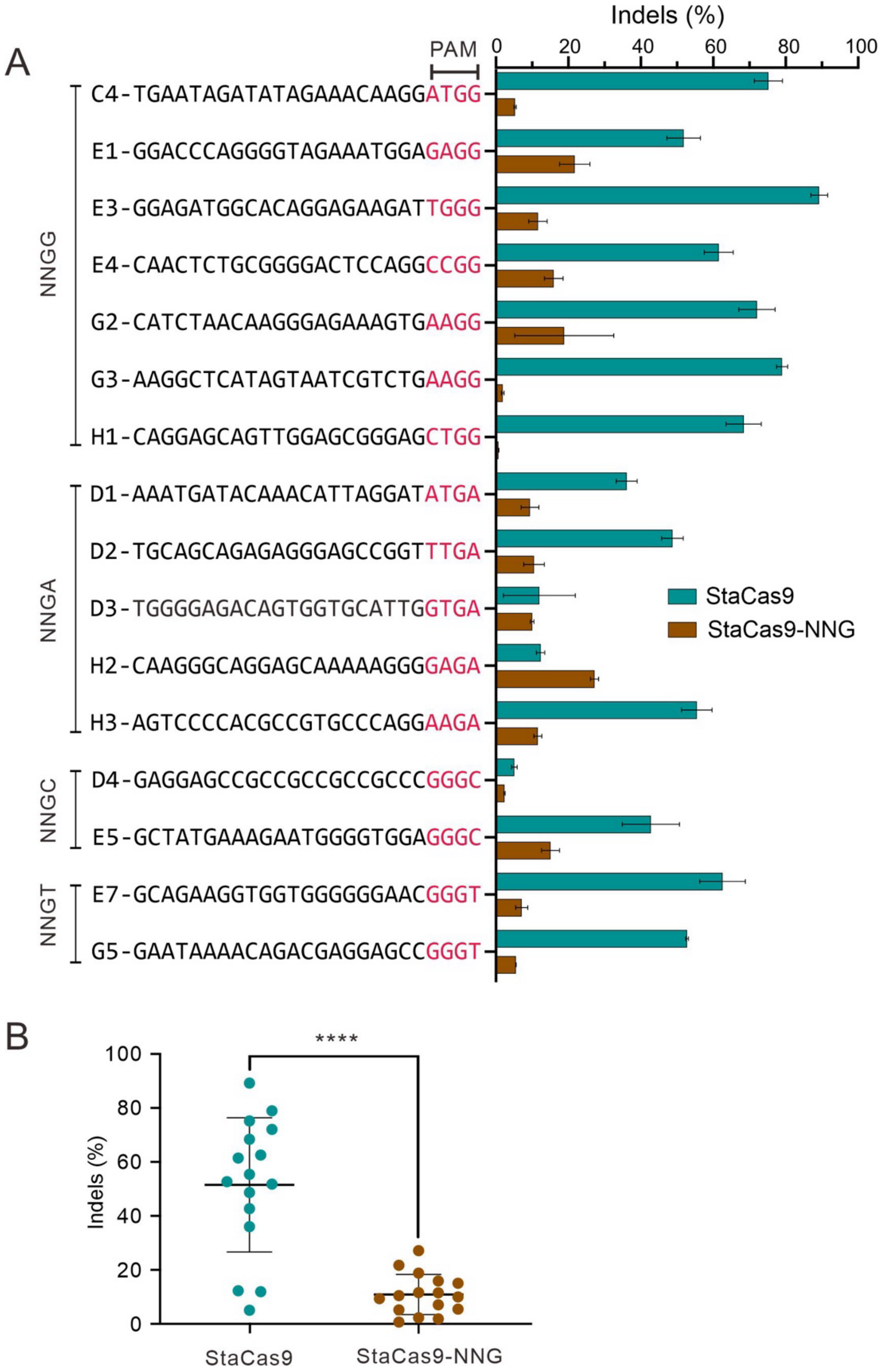


**Figure S11. Genome-editing performance of StaCas9-NNG.** (A) Genome-editing efficiencies of StaCas9-NNG and wild-type StaCas9 were evaluated at a panel of 16 endogenous genomic loci containing NNG PAMs in HEK293T cells. Data are presented as mean ± standard deviation (s.d.) from three independent experiments ($n = 3$). (B) Quantification of indel frequencies for both nucleases was determined by targeted deep sequencing. Statistical significance was assessed using a two-tailed statistical test with the following thresholds: *$P < 0.05$, **$P < 0.01$, ***$P < 0.001$, and ****$P < 0.0001$.

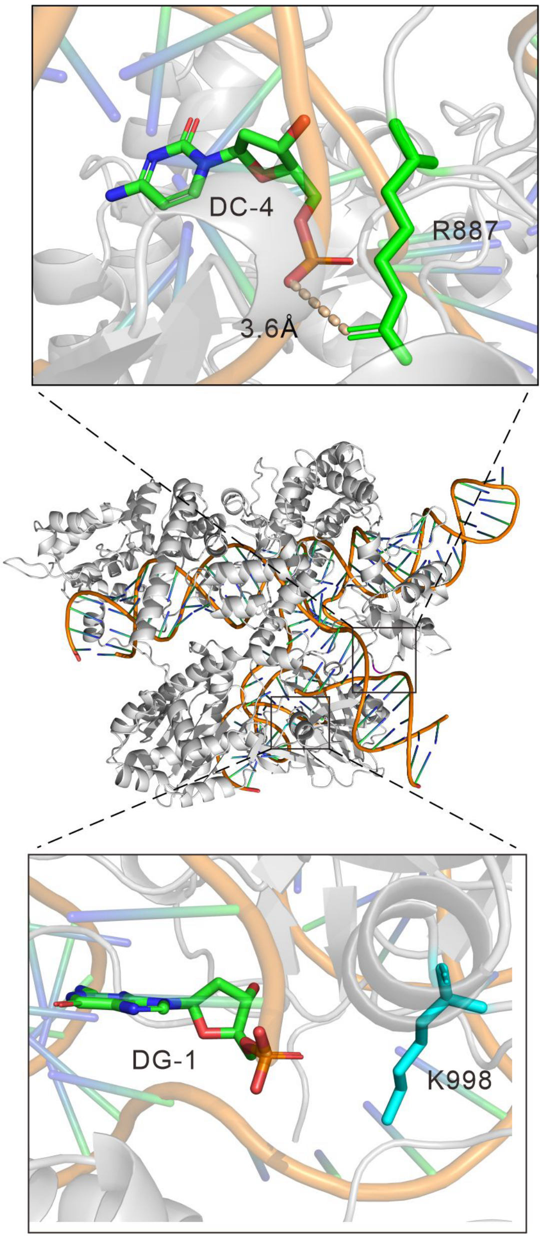


**Figure S12.** Structural basis for activity-enhancing mutations in StaCas9. Structural models of StaCas9 highlighting the positions of the activity-enhancing substitutions E887R and S998K. The left panel shows residue E887 (mutated to arginine) in proximity to the DNA backbone near the PAM-proximal region, where the introduced positive charge is predicted to strengthen electrostatic interactions with the target DNA. The right panel shows residue S998 (mutated to lysine) located adjacent to the guide RNA–DNA heteroduplex, where the substitution may enhance nucleic acid stabilization. The central panel presents an overview of StaCas9 bound to the guide RNA–DNA complex, with the locations of the engineered residues indicated. Distances are shown in angstroms (Å).
